# Targeting *EZH2* Oncogenic Splicing: Decoding the Regulatory Network and Antisense Correction

**DOI:** 10.64898/2026.01.07.697981

**Authors:** Md Rafikul Islam, Preeti Nagar, Naomi McNaughton, Shabiha Afroj Heeamoni, Md Mahbub Hasan, Shila Kandel, Panagiotis Tsakiroglou, William Brian Dalton, Omar Abdel-Wahab, Adrian R Krainer, Mohammad Alinoor Rahman

## Abstract

Recurrent mutations in splicing factors (SFs) have been established as crucial drivers of tumorigenesis in several types of blood cancer, and also common in a variety of solid tumors. Mutations change the RNA-binding preferences of SFs, promote global splicing alterations, and often generate erroneous mRNAs that are then degraded by nonsense-mediated mRNA decay (NMD). Consequently, several critical genes linked to hematopoiesis are dysregulated, leading to blood cancer. Although the field has progressed considerably in identifying aberrant genes and affected pathways, effective therapies have not yet emerged in SF-mutated cancers. To address this key gap, we instigated a gene-specific targeted strategy by unlocking the regulatory network. As a proof-of-concept, we scrutinized a tumor suppressor gene *EZH2*, which is a bona fide target in SRSF2-mutated cancer. We precisely defined splicing cis-elements in *EZH2* transcripts and illustrated the dynamic choreography of regulatory proteins in the entire splicing and NMD catalytic pathways. We uncovered a highly coordinated cross-regulation between splicing and NMD promoted by mutant SRSF2 by enhancing the deposition of critical spliceosome- and NMD-associated factors, augmenting mRNA decay to ablate tumor suppression. We then designed antisense oligonucleotides (ASOs) targeting important regulatory sites. Our lead ASO successfully corrects aberrant splicing and NMD, restores the expression and function of EZH2, and partially rescues hematopoietic defects and cellular properties. Our study demonstrates that ASO pharmacology is an actionable strategy for clinical development, challenging the existing paradigms in SF-mutated cancers.

## Introduction

The human genome preserves precise instructions for the strict regulation of gene expression. Alternative RNA splicing (AS) plays a critical role in this regulation and has received considerable attention in recent times, reflecting its extensive dysregulation in human diseases, ranging from developmental defects to deadly cancers (Black 2003; Scotti and Swanson 2016; Bradley and Anczuków 2023; Rahman et al. 2020a; Dvinge et al. 2016). Mutations in the RNA splicing factors are linked with a spectrum of human malignancies by disrupting normal RNA splicing, leading to the expression of aberrant mRNA isoform(s) that affect crucial physiological functions, promoting genomic instability, immune evasion, or altered cellular behavior (Bradley and Anczuków 2023; Yoshida et al. 2011; Lee and Abdel-Wahab 2016; Zhang et al. 2024). These alterations subsequently contribute to tumorigenesis, cancer progression, chemoresistance, and immune dysregulation.

Recurrent oncogenic heterozygous mutations in RNA splicing factors are frequent in patients with myelodysplastic syndromes (MDS), and occur in clonal hematopoiesis, acute myeloid leukemia (AML), chronic lymphocytic leukemia (CLL), and a variety of solid tumors (Bradley and Anczuków 2023; Dvinge et al. 2016; Yoshida et al. 2011; Papaemmanuil et al. 2011; Kennedy and Ebert 2017; Bejar et al. 2011; Haferlach et al. 2014). The most frequent mutations occur in SF3B1, SRSF2, U2AF1, or ZRSR2 in highly restricted residues (except for ZRSR2) (Yoshida et al. 2011; Papaemmanuil et al. 2011; Kennedy and Ebert 2017; Pellagatti et al. 2018; Thol et al. 2012). Since the discovery of splicing factor mutations(Yoshida et al. 2011), considerable progress has been made in identifying aberrantly spliced mRNAs, thanks to advancements in next-generation sequencing. Mutations in splicing factors usually change their RNA-binding properties and cause genome-wide splicing alterations (Bradley and Anczuków 2023; Dvinge et al. 2016; Zhang et al. 2024). Some of these splicing changes are directly associated with tumorigenesis (cancer drivers), while others are not linked with cancer (passenger splicing events) (Bradley and Anczuków 2023; Rahman et al. 2020a; Zhang et al. 2015; Anczuków and Krainer 2016). Cancer driver aberrant splicing events serve as important biomarkers and potential therapeutic targets.

Although initially it was predicted that mutations in different splicing factors may have common downstream effects (Yoshida et al. 2011; Haferlach et al. 2014), research in the past decade revealed that different mutations have different downstream targets. This appeared to be a critical roadblock to developing a generalized therapeutic approach to commonly treat all splicing factor-mutated cancer patients. Indeed, a generalized strategy to therapeutically target RNA splicing in cancers remains mostly ineffective to date. For example, pharmacologic inhibitors (such as E7107, H3B-8800) that target the core spliceosome component U2 small nuclear ribonucleoprotein (U2 snRNP) complex were used in preclinical studies that showed promising results with preferential killing of cancer cells in splicing factor mutant leukemias *in vivo* (Lee and Abdel-Wahab 2016; Liu et al. 2024; Seiler et al. 2018). However, the E7107-related study was terminated during clinical trial due to off-target toxicity (Hong et al. 2014; Eskens et al. 2013), whereas the H3B-8800-related study showed minimal response in clinical trial (Steensma et al. 2021). Although there has been noteworthy progress in the last 5 years, with several new US FDA-approved therapies for AML (unrelated to splicing), the 5-year survival for most AML patients remains less than 20%, and there are limited effective therapies for MDS (Daver et al. 2020; Kim et al. 2025). Thus, there remains an unmet medical need to develop more effective targeted therapeutic approaches, based on a precise understanding of molecular regulation.

Eukaryotic gene expression involves a series of highly regulated and interconnected pathways and surveillance systems to ensure its fidelity. Nonsense-mediated mRNA decay (NMD) is a post-transcriptional surveillance mechanism to selectively identify and degrade erroneous mRNAs comprising a premature termination codon (PTC). PTCs are usually generated by nonsense or frameshift mutations. AS can also generate erroneous mRNAs with a premature termination codon (PTC), which are degraded by NMD. This is termed as splicing-coupled-NMD or shortly AS-NMD. Studies in recent decades identified AS-NMD as a crucial regulator of gene expression (Nagar et al. 2023; Colombo et al. 2017; Rahman et al. 2020b). AS is regulated by coordinated interactions between splicing regulatory cis-elements in the pre-mRNA and cognate trans-acting RNA binding proteins or splicing factors (SFs) (Black 2003; Licatalosi and Darnell 2006; Cooper et al. 2009). AS regulatory cis-elements are comprised of exonic/intronic splicing enhancers (ESEs/ISEs) or silencers (ESSs/ISSs). The expression of splicing factor is often regulated in a tissue-specific or developmental stage-specific manner, resulting in differential AS outcomes to support rapidly changing biological processes (Black 2003; Licatalosi and Darnell 2006; Cooper et al. 2009; Rahman et al. 2020c). NMD is also regulated by complex interactions between mRNA and NMD-associated proteins, such as the exon junction complex (EJC) and NMD factors (UPF1, UPF2, UPF3, etc.) (Nagar et al. 2023; Lejeune et al. 2003; Popp and Maquat 2014; Metze et al. 2013; Kurosaki et al. 2019; Lykke-Andersen and Jensen 2015). The dynamic assembly of ribonucleoprotein complexes (RNPs) in the AS-NMD nexus is controlled with strict precision, which is crucial to ensure the fidelity of mRNA biogenesis. Tumor cells often differentially exploit AS-NMD for their survival benefit, such as inducing AS-NMD to downregulate tumor suppressor(s) or inhibiting AS-NMD to stabilize oncoprotein(s) (Nagar et al. 2023; Rahman et al. 2020b; Popp and Maquat 2014, 2018; Park et al. 2019). AS-NMD has been identified as a common mediator of tumorigenesis in all splicing factor-mutated cancers (Rahman et al. 2020b; DeBoever et al. 2015; Inoue et al. 2019; Kim et al. 2015; Yoshimi et al. 2019; Dolatshad et al. 2016; Zhang et al. 2019; Darman et al. 2015; Obeng et al. 2016; Dolatshad et al. 2015; Yip et al. 2017; Graubert et al. 2011; Okeyo-Owuor et al. 2015).

Understanding the molecular mechanisms and consequences of individual splicing-factor mutations is crucial for developing novel targeted splicing-based therapies that can improve the treatment of cancer. We previously showed that in addition to splicing, Pro95 hotspot mutations (P95H/L/R) in SRSF2 also affect NMD; we used a human beta globin gene (*HBB*) reporter with a β-thalassemia mutation, representing a nonsense mutation-induced mRNA decay model (NMD) (Rahman et al. 2020b). To further understand the mechanisms of AS-NMD and establish a therapeutic proof-of-concept, here we developed experimental approaches to dissect the detailed molecular mechanisms of a bona fide cancer driver AS-NMD event in *EZH2,* regulated by mutant SRSF2, and develop a molecular strategy to correct aberrant AS-NMD. Exploiting minigene reporters, model cell lines, and biochemical and functional approaches, we dissected the underlying mechanisms of aberrant inclusion of a poison exon in *EZH2* promoted by mutant SRSF2. We defined critical binding motifs of mutant SRSF2 in *EZH2* and characterized transcript-specific interacting protein assembly in the entire RNA processing pathway from splicing to NMD. Exploiting these mechanistic insights, we then developed a screening approach to modulate aberrant splicing using antisense oligonucleotides (ASO) pharmacology. The lead ASO efficiently corrected the aberrant splicing and restored the expression of EZH2 protein, evading AS-NMD. Furthermore, the ASO enhanced histone methylation (H3K27me3), improved cell viability, modulated cell cycle kinetics, reduced apoptosis, and showed promising effects in rescuing defective hematopoiesis in a preclinical model cell line derived from a leukemia patient with an SRSF2 Pro95 mutation. Our approach is promising and will pave the way for future targeted therapeutic development in splicing factor-mutated cancer.

## Results

### Aberrant splicing of the *EZH2* poison exon is a crucial tumorigenic target in SRSF2-mutated blood cancer

We previously investigated AML patient RNA-sequencing data set from the Cancer Genome Atlas (TCGA-LAML) for SRSF2 Pro95 mutant (SRSF2^Mut^, n=6) versus SRSF2 wild-type (SRSF2^WT^, n=109) (Rahman et al. 2020b). We have reanalyzed the data to detect a potential tumorigenic target to push our study forward towards therapeutic development dependent on mechanistic insights. Among 843 differential splicing events identified in our analysis, about half of the events are cassette exon splicing (ES), and the remaining half include alternative splice site selection (ASS) and intron retention (IR) events in similar proportions (**Fig. 1A**). One striking observation is that 135 mRNA isoforms (∼16%) among these events are NMD targets (AS-NMD), which are distributed among all types of splicing events, but with higher representation in ES group (**Fig. 1B** and **C**). Aberrant AS-NMD has also been identified as a major mechanism of tumorigenesis in SF3B1- and U2AF1-mutated patients (Rahman et al. 2020b; DeBoever et al. 2015; Inoue et al. 2019; Kim et al. 2015; Yoshimi et al. 2019; Dolatshad et al. 2016; Zhang et al. 2019; Darman et al. 2015; Obeng et al. 2016; Dolatshad et al. 2015; Yip et al. 2017; Graubert et al. 2011; Okeyo-Owuor et al. 2015). Analysis of cassette exons (SRSF2^Mut^ vs. SRSF2^WT^) to quantify enriched motifs using the MEME (multiple expectation maximization for motif elicitation) (Bailey et al. 2009) shows preferential enrichment of (G/C)C(A/T)G motifs in promoted exons, and GG(A/T)G motifs in repressed exons (**Fig. 1D** and **E**). These consensus motifs retrieved from RNA-sequencing is consistent with previous studies, including samples from cancer patients, animal models, and cell lines (Zhang et al. 2015; Kim et al. 2015; Komeno et al. 2015). A recent study determined motif enrichment from *in vivo* HITS-CLIP (high-throughput sequencing of RNA isolated by crosslinking immunoprecipitation) and also identified similar motifs (Liang et al. 2018). Taken together, these analyses show that CCWG is the consensus motif preferred for SRSF2^Mut^ (W=A/T), whereas GGWG is the motif for SRSF2^WT^. The change in RNA binding preferences causes a gain of function in SRSF2^Mut^ and triggers aberrant splicing of a set of target genes, some of which are strongly associated with defective hematopoietic differentiation (Zhang et al. 2015; Kim et al. 2015; Yoshimi et al. 2019; Komeno et al. 2015; Liang et al. 2018). Among consistently identified genes in multiple studies, a few validated and noteworthy cancer-relevant aberrant splicing targets include *EZH2, INTS3, and CLK3* (Kim et al. 2025; Rahman et al. 2020b; Kim et al. 2015; Yoshimi et al. 2019). *EZH2* encodes the enhancer of zeste homolog 2 protein (EZH2), which catalyzes methylation of histone H3 at lysine 27 (H3K27me3), and therefore functions in chromatin remodeling (Tan et al. 2014) (**Fig. 1F**). EZH2, as a component of the polycomb repressive complex 2 (PRC2), was reported to suppress gene expression and regulate cell fate (Tan et al. 2014). A poison exon (PE) in *EZH2* is exclusively included in SRSF2^Mut^ patients (**Fig. 1G)**, which generates a PTC and triggers the degradation of *EZH2* transcripts by AS-NMD (Kim et al. 2015; Yoshimi et al. 2019; Rahman et al. 2020b). Downregulation of EZH2 protein expression through AS-NMD consequently leads to defective hematopoietic differentiation in SRSF2^Mut^ patients (Kim et al. 2015; Yoshimi et al. 2019; Rahman et al. 2020b).

**Figure 1.**
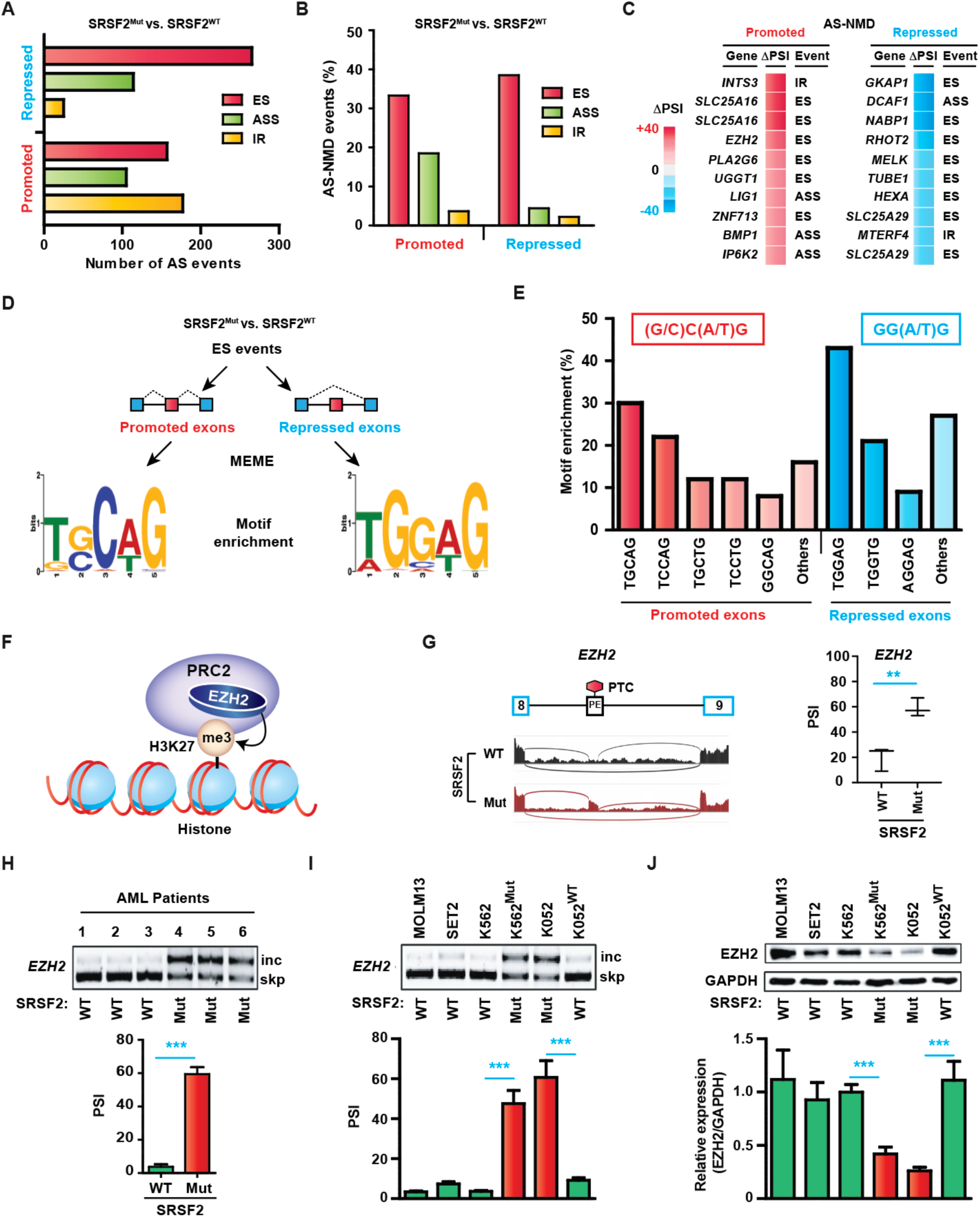
Tumorigenic aberrant splicing targets in SRSF2-mutated blood cancer. **(A)** Differential alternative splicing (AS) events in acute myeloid leukemia (AML) patients with heterozygous oncogenic mutations at Pro95 of SRSF2 (SRSF2^Mut^) compared to wild-type SRSF2 (SRSF2^WT^), derived from TCGA-LAML RNA-seq data analysis (Rahman et al. 2020b). ASS: alternative splice site; ES: cassette exon; IR: intron retention. **(B)** Distribution of NMD targets (AS-NMD) from panel **(A)** among different types of altered splicing events. **(C)** Top ten AS-NMD targets are enlisted in promoted and repressed alternative splicing events, respectively. ι1PSI: changes in percent spliced-in (SRSF2^Mut^ vs. SRSF2^WT^). **(D)** Schematic of motif enrichment analysis in promoted or repressed cassette exons (SRSF2^Mut^ vs. SRSF2^WT^) using MEME. **(E)** Proportion of different motifs enriched in promoted or repressed cassette exons exons. Consensus sequences of enriched motifs are shown above in enclosed boxes. **(F)** Schematic of EZH2 protein function in histone methylation (H327Kme3) and chromatin regulation. PRC2: polycomb repressing complex. **(G)** Left, representative Sashimi plots showing *EZH2* poison exon inclusion in AML samples from the genotypes indicated. Right, quantification of *EZH2* poison exon inclusion as percent spliced-in (PSI) in AML samples from the genotypes indicated (mean ± SD, n=3, **p<0.001, t test). **(H)** RT-PCR gel showing *EZH2* poison exon inclusion (inc) and skipping (skp) in an independent set of AML samples with SRSF2^WT^ and SRSF2^Mut^. Quantification of *EZH2* poison exon inclusion as PSI is shown by a bar graph below the gel (mean ± SD, n=3, ***p<0.001, t test) **(I)** Representative RT-PCR gel showing *EZH2* poison exon inclusion and skipping in different human leukemia cell lines from the genotypes indicated below. Quantification of *EZH2* poison exon inclusion as PSI is shown by a bar graph below the gel (mean ± SD, n=3, ***p<0.001, t test). **(J)** Western blotting of EZH2 protein expression in different human leukemia cell lines from panel **(I)**. GAPDH was used as an internal control. Relative quantification of EZH2 protein expression (EZH2/GAPDH) is shown by a bar graph below the gel (mean ± SD, n=3, ***p<0.001, t test).

We confirmed aberrant poison exon inclusion in *EZH2* in an independent pool of AML patient RNA samples with SRSF2^Mut^ compared to SRSF2^WT^ (**Fig. 1H)**. This was also consistently shown in several other studies (Kim et al. 2015; Yoshimi et al. 2019; Grimm et al. 2021; Masaki et al. 2019). We analyzed several cell lines originating from leukemia patients, with inherently harboring wild-type (SRSF2^WT/WT^) or heterozygous mutant (SRSF2^Mut/WT^) genotypes. These included MOLM-13 (SRSF2^WT/WT^), SET2 (SRSF2^WT/WT^), K562 (SRSF2^WT/WT^), and K052 (SRSF2^P95H/WT^), where SRSF2 genotypes are mentioned in brackets. RT-PCR and western blotting showed exclusive poison exon inclusion into *EZH2* mRNA and concomitant protein downregulation only in K052 (SRSF2^P95H/WT^), but not in the other cell lines (**Fig. 1I** and **J**). We additionally included an engineered K562 cell line, termed as K562^Mut^, in which the *SRSF2* P95H mutation was introduced into the endogenous locus as a heterozygous mutation (SRSF2^P95H/WT^). As expected, K562^Mut^ showed enhanced poison exon inclusion into *EZH2* mRNA compared to K562 (**Fig. 1I**). This was also reflected at the protein level, with downregulation of EZH2 protein expression (**Fig. 1J**). Finally, we included another engineered K052 cell line, termed as K052^WT^, in which the *SRSF2* P95H mutation was reverted to *SRSF2* WT in the endogenous gene locus (SRSF2^WT/WT^). Indeed, K052^WT^ showed a reduction of poison exon inclusion into *EZH2* mRNA, and upregulated EZH2 protein expression compared to K052 (SRSF2^P95H/WT^) (**Fig. 1I** and **J**). These rigorous experimental results clearly confirmed the SRSF2^Mut^-dependent poison exon inclusion activity in *EZH2*. We, therefore, selected *EZH2* as a bona fide target for downstream mechanistic analyses aimed at developing a potential therapeutic strategy.

### Experimental models to characterize mechanisms of regulation by mutant SRSF2

To study the precise mechanisms of action that link mutant SRSF2 to tumorigenesis, we engineered K562 cells and stably integrated a doxycycline (DOX)-inducible construct to conditionally express cDNA encoding FLAG-tagged SRSF2^WT^ or SRSF2^P95H^ (**Fig. 2A**). These cell lines are termed K562(DOX)SRSF2^WT^ and K562(DOX)SRSF2^P95H^, respectively. Since mutations in SRSF2 are frequently identified in hematopoietic tissues, as a control to examine tissue-specific regulation, we also engineered a non-hematopoietic HeLa cell line expressing DOX-inducible FLAG-tagged SRSF2^WT^ or SRSF2^P95H^ to examine any tissue-specific regulation. These cell lines allow the study of SRSF2^WT^- or SRSF2^P95H^-specific regulatory mechanisms in splicing and NMD, following DOX-inducible expression. Although SRSF2 is involved in multiple biological pathways, conditional expression over a short period is intended to exclude indirect or feedback effects from other pathways, and/or long-term cellular effects exerted by the mutant factor.

**Figure 2.**
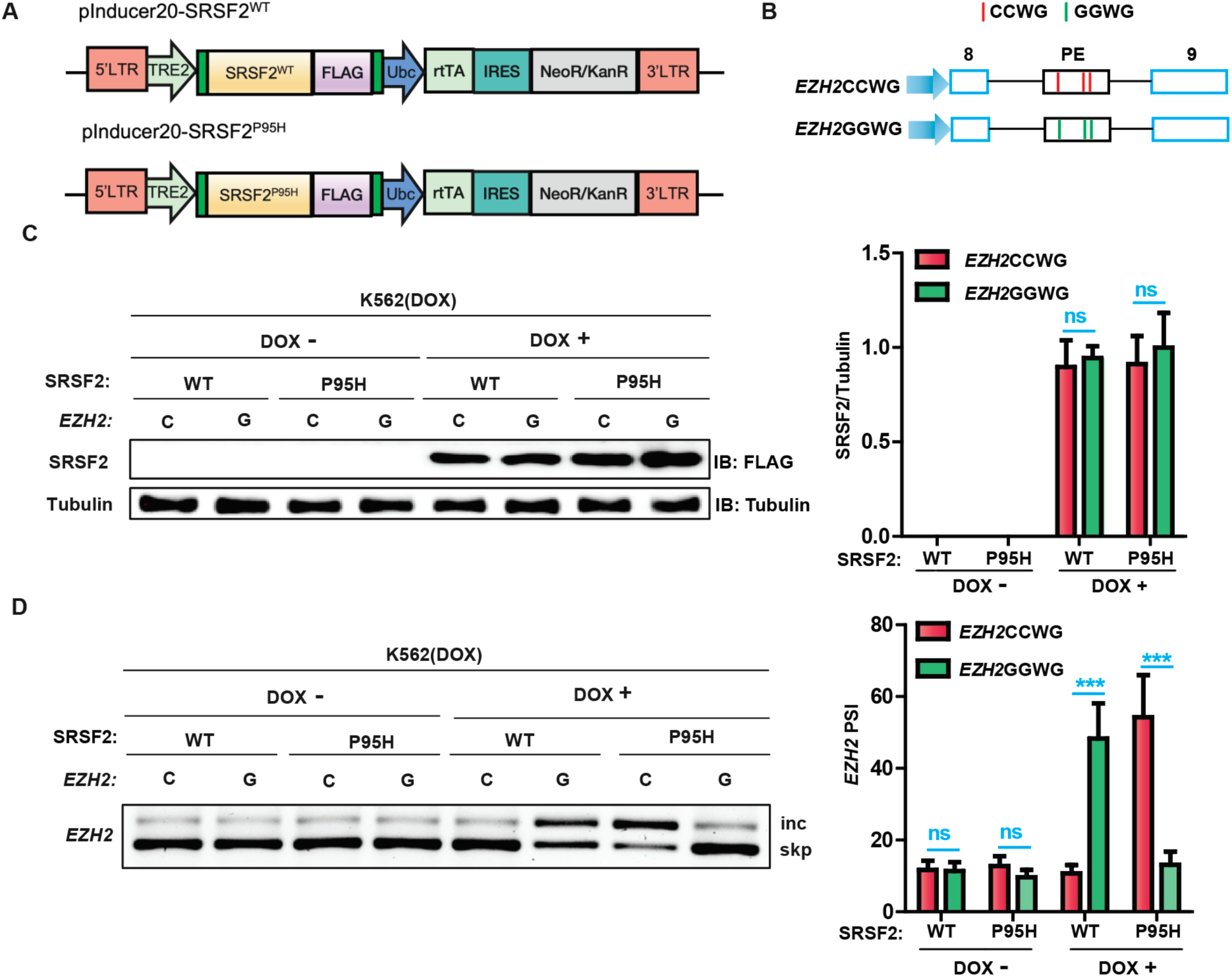
Alternative splicing of the *EZH2* poison exon is regulated by the presence or absence of specific ESE elements. **(A)** Diagram of DOX-inducible FLAG-tagged cDNA constructs encoding SRSF2 wild-type (SRSF2^WT^) or Pro95 mutant (SRSF2^P95H^) to generate K562(DOX) stable cell lines expressing either SRSF2^WT^ or SRSF2^P95H^ in the presence of DOX. **(B)** Diagram (not to scale) of *EZH2* minigene reporters harboring different combinations of SRSF2 (Mut or WT) motifs within the poison exon. *EZH2*CCWG (C) contains native gene sequences with three SRSF2^Mut^-binding motifs (CCWG), whereas *EZH2*GGWG (G) contains three SRSF2^WT^-binding motifs (GGWG) replacing CCWG motifs (W=A/U). **(C)** Left, western blotting (WB) of the indicated K562(DOX) cells co-transfected with the indicated *EZH2* minigene reporters with the indicated antibodies. Tubulin was used as an internal control. Right, quantification of SRSF2 (WT or P95H) protein expression (normalized against tubulin) is shown by a bar graph (mean ± SD, n=3, ns: not significant, t test). **(D)** Left, representative RT-PCR gel showing *EZH2* minigene reporter-specific poison exon inclusion (inc) and skipping (skp) in the indicated K562(DOX) cells co-transfected with the indicated *EZH2* minigene reporters. Right, quantification of *EZH2* minigene reporter-specific poison exon inclusion as percent spliced-in (PSI) is shown by a bar graph (mean ± SD, n=3, ***p<0.001, ns: not significant, t test).

### *EZH2* PE splicing is regulated by the presence or absence of a specific exonic splicing enhancer

Inclusion of an alternative exon is usually regulated by an exonic splicing enhancer (ESE), and often by an intronic splicing enhancer (ISE) (Licatalosi and Darnell 2006; Black 2003). As noted earlier, Pro95 mutations in SRSF2 change its RNA-binding preference from GGWG to CCWG. Tabulating these motifs revealed that *EZH2* poison exon harbors three preferred binding motifs (CCWG) of SRSF2^Mut^, but no binding motif (GGWG) of SRSF2^WT^. Splicing regulatory ESE in an alternative exon often harbors multiple cis-elements or binding motifs, and these motifs coordinately regulate the binding of splicing factor(s) and subsequent splicing modulation (Feng et al. 2019; Nasrin et al. 2014, 2025). To systematically identify splicing regulatory elements in *EZH2* poison exon ESE, we employed an *EZH2* minigene reporter (*EZH2*CCWG) with native gene sequences(Kim et al. 2015), consisting of three binding motifs of SRSF2^Mut^ in the PE (**Fig. 2B**). We also employed another modified minigene (*EZH2*GGWG), where all of the three binding motifs of mutant SRSF2 were changed to preferred motifs for SRSF2^WT^ (**Fig. 2B**). We then transfected these minigene reporters into DOX-inducible K562 cells for splicing analyses. We first confirmed the comparable levels of expression of SRSF2^WT^ and SRSF2^Mut^ upon DOX treatment by western blotting (**Fig. 2C**). We then performed RT-PCR to study minigene reporter-specific splicing profiles by employing a reporter-specific primer during cDNA synthesis. In the absence of DOX, we found no changes in *EZH2* PE splicing between SRSF2^P95H^ and SRSF2^WT^ expressing cells (**Fig. 2D**). To our expectation, DOX treatment could clearly enhance PE inclusion in *EZH2*CCWG reporter in SRSF2^P95H^ expressing cells, but not in SRSF2^WT^ expressing cells (**Fig. 2D**). These data clearly demonstrated CCWG-dependent ESE-activity of SRSF2^Mut^ in native *EZH2* reporter (*EZH2*CCWG). Surprisingly, DOX treatment could also enhance PE inclusion in the *EZH2*GGWG reporter in SRSF2^WT^ expressing cells, but not in SRSF2^P95H^ expressing cells. These data suggest that the lack of SRSF2^WT^-binding motif in *EZH2* PE disables exon inclusion in normal cells expressing SRSF2^WT^, whereas the presence of the SRSF2^Mut^ motif promotes *EZH2* PE inclusion in SRSF2-mutated blood cancer cells. We also performed the experiment in DOX-inducible HeLa cells. Our data in HeLa cells showed consistent results with that of K562 cells (**Fig. 2**), thereby ruling out any tissue-specific differential regulation. Taken together, we can conclude that the presence or absence of preferred RNA-binding motif(s) in the *EZH2* PE ESE can dictate the splicing consequences promoted by SRSF2^Mut^ or SRSF2^WT^.

### Systematic identification of a crucial functional motif in the ESE of *EZH2* PE

Among multiple motifs in a splicing regulatory ESE, a specific motif often plays a crucial role in cooperative binding or stable association of splicing factor(s) with RNA (Feng et al. 2019; Nasrin et al. 2014, 2025). Therefore, identification of a crucial functional motif could provide the opportunity to develop a splicing modulation strategy by targeting a specific region of RNA either by generating steric hindrance, affecting RNA structure, or interfering with the binding of splicing factor(s). To this end, we wanted to define the contribution of individual binding motifs on SRSF2^Mut^-mediated *EZH2* PE inclusion. We employed systematic inactivation of individual binding motifs in the minigene reporter, followed by analyzing splicing consequences. To inactivate the SRSF2^Mut^ binding motif, we changed the CCWG motif to the AAWG motif, which is preferred neither by SRSF2^Mut^ nor by SRSF2^WT^, as we showed previously(Rahman et al. 2020b). Note that among the three binding motifs of SRSF2^Mut^ in the *EZH2* poison exon, motifs 2 and 3 overlap. Therefore, we employed three mutant versions of *EZH2* reporter: *EZH2*MUT1 harbors inactivated motif 1; *EZH2*MUT2 harbors inactivated motifs 2 and 3; and *EZH2*MUT3 harbors inactivated motifs 1, 2, and 3 (**Fig. 3A**). We then performed minigene reporter-specific splicing assay using RT-PCR. In K562(DOX)SRSF2^WT^ or K562(DOX)SRSF2^P95H^, in the absence of DOX, we observed no considerable changes in *EZH2* PE inclusion (**Fig. 3B**). However, upon DOX treatment in K562(DOX)SRSF2^P95H^, we found significant loss of *EZH2* PE inclusion in *EZH2*MUT1 and *EZH2*MUT3, compared to *EZH2*CCWG (Native) (**Fig. 3B**). Note that motif 1 was inactivated in both *EZH2*MUT1 and *EZH2*MUT3. Therefore, the results highlight the importance of motif 1 for SRSF2^Mut^ and also suggest that motifs 2 and 3 are not enough in the absence of motif 1. In contrast, we observed a slight loss of *EZH2* PE inclusion in *EZH2*MUT2 compared to *EZH2*CCWG (Native) (**Fig. 3B**). This observation indicates that motif 1 is functional even in the absence of motif 2 and motif 3; however, motif 2 and motif 3 have additive effects, which may function in cooperation with motif 1. It is important to note that the above changes were not evident upon DOX induction in K562(DOX)SRSF2^WT^, as expected due to the lack of SRSF2^WT^ binding motif (**Fig. 3B**). Taken together, these data clearly suggest the essentiality and critical importance of motif1 in regulating SRSF2^Mut^-mediated *EZH2* PE inclusion.

**Figure 3.**
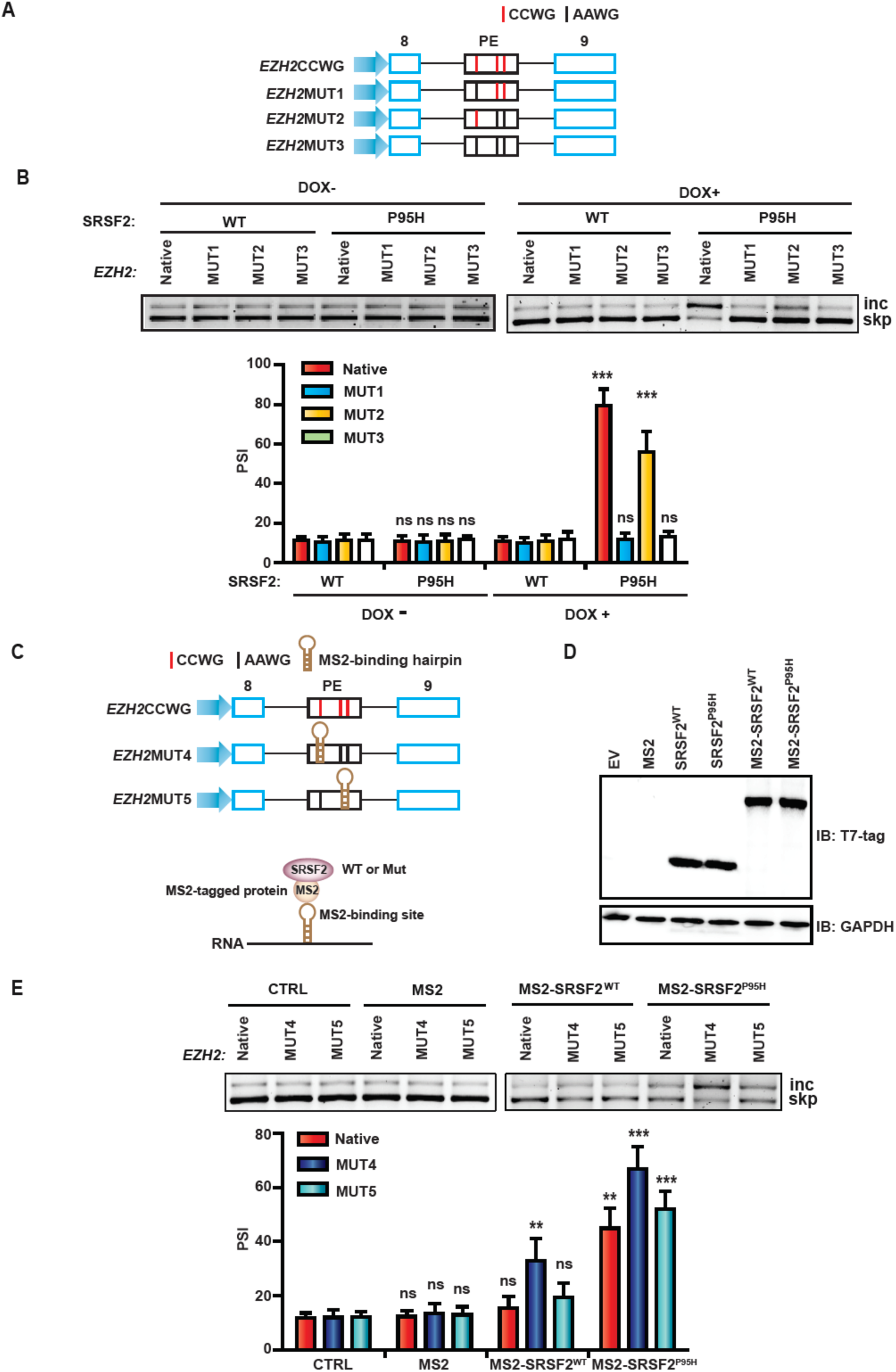
Decoding crucial ESE motif in the *EZH2* poison exon regulated by SRSF2^Mut^. **(A)** Diagram (not to scale) of *EZH2*CCWG (Native) and three mutant minigene reporters. *EZH2*CCWG (Native) contains native gene sequences with three SRSF2^Mut^-binding motifs (CCWG): motif 1, motif 2, and motif 3, respectively (W=A/U). These CCWG motifs were mutated to AAWG to inactivate the binding of SRSF2^Mut^ in different combinations in mutant reporters. **(B)** Top, representative RT-PCR gels showing *EZH2* minigene reporter-specific poison exon inclusion (inc) and skipping (skp) in K562(DOX)SRSF2^WT^ or K562(DOX)SRSF2^P95H^ cells co-transfected with the indicated *EZH2* minigene reporters. Bottom, quantification of *EZH2* minigene reporter-specific poison exon inclusion as percent spliced-in (PSI) is shown by a bar graph (mean ± SD, n=3, ***p<0.001, ns: not significant, t test, SRSF2^P95H^ vs. SRSF2^WT^). **(C)** Top, diagram (not to scale) of *EZH2*CCWG (Native) and two mutant minigene reporters. In *EZH2*MUT4, motif 1 was replaced by the MS2 coat protein-binding hairpin, and motifs 2 and 3 were inactivated to AAWG. In *EZH2*MUT5, motif 2 and motif 3 were replaced by the MS2 coat protein-binding hairpin, and motif 1 was inactivated to AAWG. Bottom, schematic of targeted recruitment of MS2-tagged SRSF2 (WT or Mut) to a specific site of a transcript harboring the MS2 coat protein-binding hairpin. **(D)** Immunoblotting (IB) of K562 cells transfected with the indicated cDNA vectors. Note that the empty vector (CTRL) and only MS2 (MS2) have no T7-tag. **(E)** Top, representative RT-PCR gels showing *EZH2* minigene reporter-specific poison exon inclusion (inc) and skipping (skp) in K562 cells co-transfected with the indicated *EZH2* minigene reporters and cDNAs. Bottom, quantification of *EZH2* minigene reporter-specific poison exon inclusion as percent spliced-in (PSI) is shown by a bar graph (mean ± SD, n=3, **p<0.01, ***p<0.001, ns: not significant, t test, compared to corresponding CTRL).

### MS2-mediated tethering of SRSF2 to a specific binding motif in *EZH2* PE is sufficient to promote exon inclusion

Having identified the functional motifs for ESE activity in *EZH2* poison exon, we next sought to identify whether site-specific tethering of SRSF2^Mut^ to the motif site could reproduce ESE-activity (**Fig. 3C**). To this end, we generated two additional versions of *EZH2* reporter: *EZH2*MUT4, where we replaced motif 1 with the MS2 coat protein-binding hairpin sequences (Rahman et al. 2020b), and inactivated both motif 2 and 3 to AAWG; and *EZH2*MUT5, where we replaced both motif 2 and motif 3 with the MS2 coat protein-binding hairpin sequences, and inactivated motif 1 to AAWG (**Fig. 3C**). These reporters, therefore, allow to evaluate the effect of MS2-mediated artificial tethering of SRSF2 for only one site at a time. We generated MS2-tagged-T7-SRSF2^WT^ (MS2-SRSF2^WT^) and MS2-tagged-T7-SRSF2^Mut^ (MS2-SRSF2^P95H^) cDNA constructs and confirmed their comparable expression in K562 (**Fig. 3D**). We then simultaneously transfected K562 cells with an *EZH2* reporter and MS2-tagged SRSF2 cDNA (WT or Mut) and performed minigene reporter-specific splicing assay using RT-PCR. Recruitment of MS2-SRSF2^Mut^ to only motif 1 site (*EZH2*MUT4) significantly enhanced exon inclusion activity (**Fig. 3E**). In contrast, recruitment of MS2-SRSF2^Mut^ to the site spanning motif 2 and motif 3 (*EZH2*MUT5) also enhanced exon inclusion activity, but to a lesser extent than only motif 1 site (*EZH2*MUT4) (**Fig. 3E**). We also observed exon inclusion activity of MS2-SRSF2^Mut^ in *EZH*CCWG (Native) reporter without MS2 hairpin, because MS2-SRSF2^Mut^ can still bind to CCWG motifs through its SRSF2^Mut^ part of the fusion protein. Note that MS2 alone showed no effects in any of the reporters, ruling out any possible experimental artifacts promoted by the MS2-tag. Interestingly, recruitment of MS2-SRSF2^WT^ to only motif 1 site also promoted exon inclusion activity, because MS2-SRSF2^WT^ was able to bind the PE through MS2 hairpin even in the absence of GGWG motif (**Fig. 3E**). These data again support that the presence or absence of preferred RNA-binding motif(s) in an ESE of a targeted exon could dictate the splicing consequences promoted by SRSF2^Mut^ or SRSF2^WT^.

### Mutant SRSF2 enhances the deposition of several key spliceosome components and NMD-associated factors to augment AS-NMD

AS-NMD is regulated at multiple levels of RNA metabolism, including splicing of pre-mRNA, mRNA surveillance to selectively identify erroneous mRNAs with a premature termination codon, mRNA transport, and rapid degradation of faulty mRNAs. These metabolisms are choreographed with dynamic interactions between RNA and ribonucleoproteins (RNPs) at individual stages (Nagar et al. 2023; Kurosaki et al. 2019; Lykke-Andersen and Jensen 2015; Woodward et al. 2017). RNA-binding proteins or splicing factors often positively or negatively regulate assembly of RNPs, and differentially regulate AS-NMD, therefore function as molecular regulators of gene expression (Nagar et al. 2023; Rahman et al. 2020b). To identify how assembly of RNPs is regulated by SRSF2^Mut^ in *EZH2* transcripts, we exploited the MS2 coat protein-mediated RNPs pull-down system that we described before (Rahman et al. 2020b). We constructed two additional modified versions of *EZH2 reporters, EZH2*CCWG-3×MS2 *and EZH2*AAWG-3×MS2, where we introduced three MS2-binding hairpin sequences at the 3’ end of the reporters (**Fig. 4A**). Transcripts (both pre-mRNA and mRNA) originating from these minigenes will generate MS2-binding hairpins. After transfecting these reporters in K562(DOX)SRSF2^WT^ or K562(DOX)SRSF2^P95H^ cells, MS2 hairpins-containing transcripts and associated RNPs can be selectively captured using recombinant MS2-tagged maltose binding protein (MS2-MBP) and amylose resin beads (**Fig. 4B**). Purified RNPs can then be resolved and quantified by western blotting. We performed the experiments and quantified the assembly of major spliceosome components and NMD-associated proteins, such as exon junction complex (EJC) and NMD factors into *EZH2* transcripts (**Fig. 4C** and **Fig. 5**). Our results showed that SRSF2^P95H^ expression (upon DOX treatment) enhances the assembly of several key spliceosome components in *EZH2*CCWG-3xMS2 transcripts compared to *EZH2*AAWG-3xMS2, including U1snRNP 70K (U1-70K), U1 snRNP A (U1A), U1 snRNP C (U1C), U2AF65, and U2AF35, but not SF3B1 (**Fig. 5A**). Similarly, SRSF2^P95H^ expression enhances deposition of several EJC components, such as eIF4A3, Y14, MAGOH, but not MLN51 (**Fig. 5B**). Among NMD factors, SRSF2^P95H^ expression enhances deposition of UPF1, UPF2, UPF3 in *EZH2*CCWG-3xMS2 transcripts compared to *EZH2*AAWG-3xMS2 (**Fig. 5B**). These enhancements were not observed in K562(DOX)SRSF2^P95H^ cells without DOX treatment. These results clearly pointed out that SRSF2^Mut^-regulated augmented AS-NMD activity is dependent on sequence-specific binding to *EZH*2 PE RNA.

**Figure 4.**
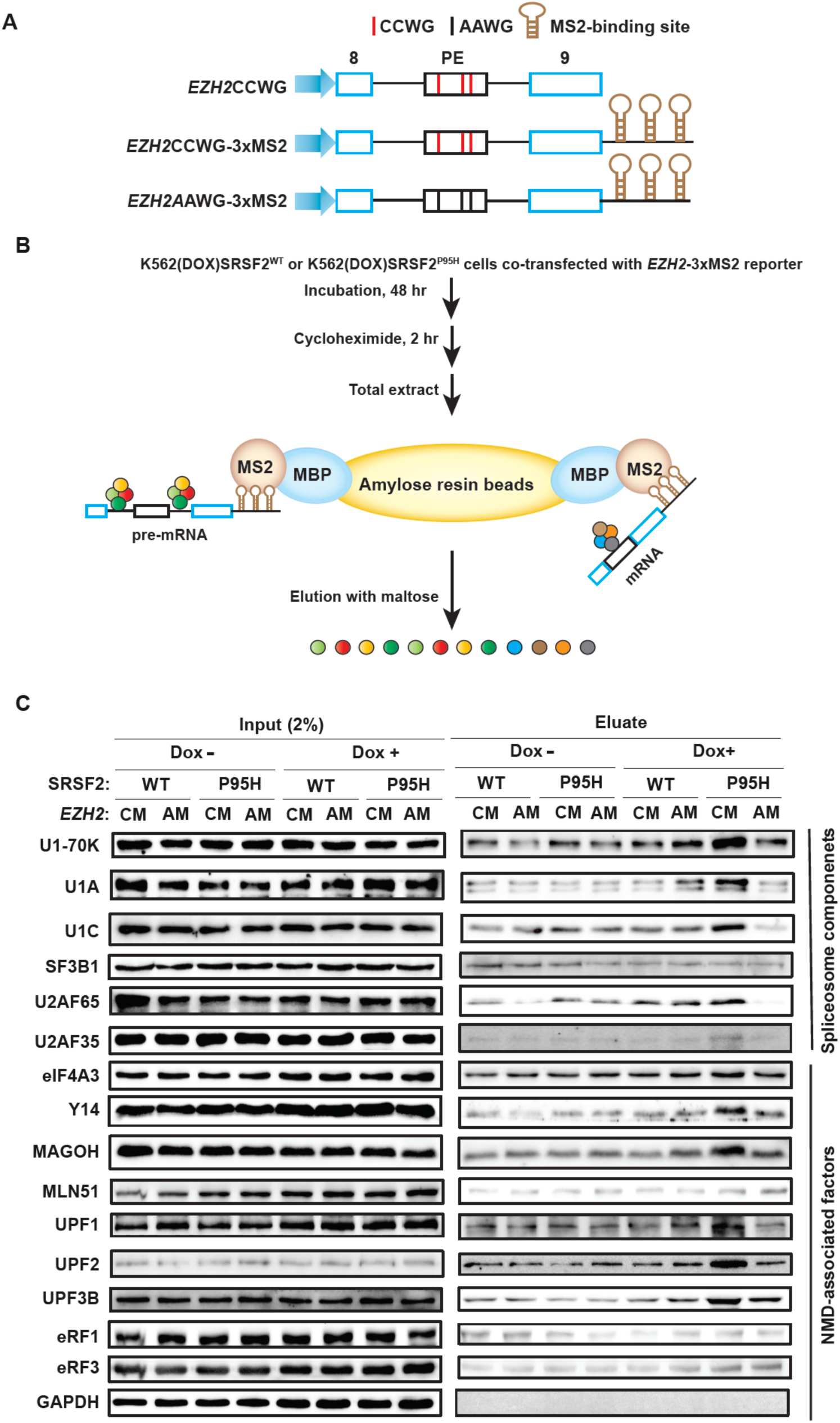
Purification of ribonucleoproteins (RNPs) from *EZH2* minigene reporter-specific transcripts. **(A)** Diagram (not to scale) of *EZH2-*3xMS2 minigene reporters harboring three MS2 coat protein-binding hairpin sequences downstream of exon 9. *EZH2*CCWG*-*3xMS2 (CM) contains native gene sequences with three SRSF2^Mut^-binding motifs (CCWG). *EZH2*AAWG*-*3xMS2 (AM) contains three AAWG motifs (not a binding site for SRSF2^Mut^ or SRSF2^WT^) replacing respective CCWG motifs. **(B)** Schematics of MS2-mediated RNPs purification from *EZH2-*3xMS2-specific transcripts (both pre-mRNA and mRNA). Recombinant MS2-tagged maltose-binding protein (MS2-MBP) and amylose resin beads were used to purify the MS2 hairpin-containing transcripts originating from *EZH2-*3xMS2 reporter transfected into K562(DOX)SRSF^WT^ or K562(DOX)SRSF2^P95H^ cells. **(C)** Representative western blotting of input or purified RNPs from the indicated K562(DOX) cells co-transfected with the indicated reporters.

**Figure 5.**
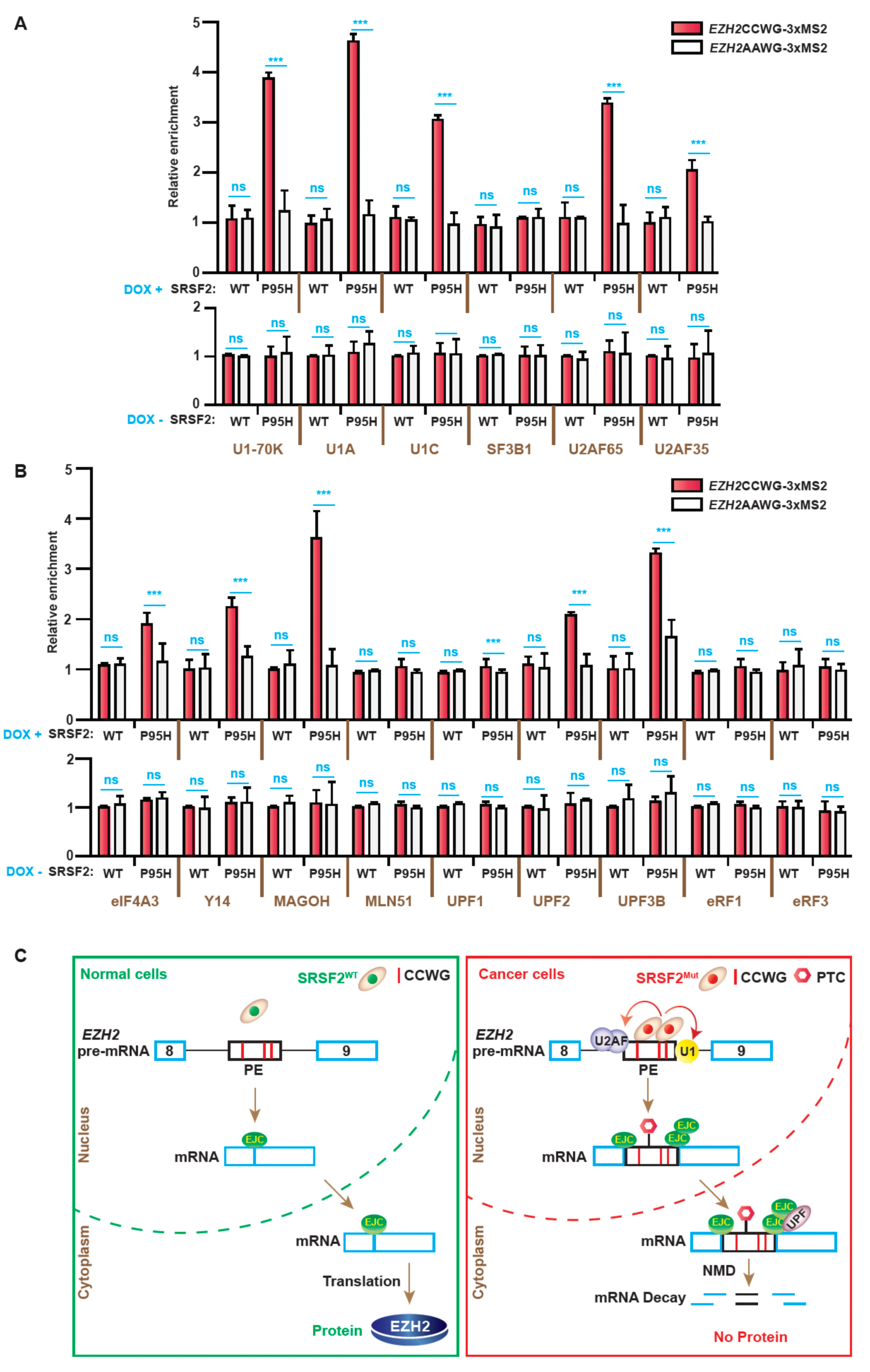
SRSF2^Mut^ elicits AS-NMD of *EZH2* by enhancing deposition of several key spliceosome components and NMD-associated factors. **(A)** Quantification of the relative enrichment of spliceosome components into *EZH2*CCWG*-*3xMS2 (CM) transcripts compared to *EZH2*AAWG*-*3xMS2 (AM) transcripts purified from transfected K562(DOX) cells with the indicated genotypes in the presence or absence of DOX (mean ± SD, n=3, ***p<0.001, ns: not significant, t test). **(B)** Quantification of the relative enrichment of EJC components and NMD factors into *EZH2*CCWG*-*3xMS2 transcripts compared to *EZH2*AAWG*-*3xMS2 transcripts purified from transfected K562(DOX) cells with the indicated genotypes in the presence or absence of DOX (mean ± SD, n=3, ***p<0.001, ns: not significant, t test). **(C)** Schematics of AS-NMD regulation of *EZH2* in normal cells (SRSF2^WT^) and SRSF2 Pro95 mutated cancer cells (SRSF2^Mut^). Dynamic interactions of key spliceosome components, EJC proteins, and NMD factors with *EZH2* transcripts derived from panels **(A)** and **(B)** are shown. PTC: premature termination codon.

In contrast, we didn’t observe any enhancements of RNPs deposition for SRSF2^WT^ expression (upon DOX treatment) in *EZH2*CCW*G*-3xMS2 transcripts compared to *EZH2*AAWG-3xMS2 (**Fig. 5A and B**). This is consistent with our expectation due to the lack of preferred binding motifs of SRSF2^WT^ in *EZH2*CCW*G*-3xMS2, therefore, a lack of AS-NMD activity.

Taken together, our data propose that sequence-specific binding of SRSF2^Mut^ into *EZH2* PE enhances spliceosome assembly in the flanking splice sites to promote exon inclusion, which is followed by enhanced EJC deposition and subsequent recruitment of NMD factors to facilitate AS-NMD (**Fig. 5C**).

### Screening of antisense oligonucleotides to modulate aberrant splicing of *EZH2* poison exon using minigene reporter model

We next wanted to extend our study to develop a molecular strategy to suppress the aberrant inclusion of the *EZH2* poison exon to restore the expression of the EZH2 protein. In the past decade, several pharmacological small-molecule compounds were employed to modulate aberrant splicing in cancer (Lee and Abdel-Wahab 2016; Liu et al. 2024; Seiler et al. 2018). Although a few progressed to clinical trials, they were terminated due to off-target effects (Hong et al. 2014; Eskens et al. 2013) or appeared ineffective (Steensma et al. 2021). Therefore, we undertook a gene-specific targeted approach to correct aberrant splicing of cancer-driver gene(s). We chose antisense pharmacology. Antisense oligonucleotides (ASO) are short synthetic single-stranded nucleic acid molecules that are usually designed to bind to complementary sequences on a target RNA (Rigo et al. 2012; Havens and Hastings 2016). ASO may inhibit the binding of splicing factor(s) or RNA-binding protein(s) or generate steric hindrance to interfere with spliceosome assembly, resulting in differential alternative splicing outcomes. The efficacy of ASO depends on multiple factors (Havens and Hastings 2016; Rigo et al. 2012), such as target site or sequence, binding affinity, cellular delivery, potency, resistance to nuclease, stability, etc. Based on our detail mechanistic characterization of *EZH2* splicing as described above (**Fig. 5C**), we designed four ASOs: ASO1 targeting the flanking 3’ splice site of the PE; ASO2 targeting the SRSF2^Mut^ binding site spanning motif 1 in the PE; ASO3 targeting the SRSF2^Mut^ binding sites spanning both motif 2 & motif 3 in the PE; and ASO4 targeting the flanking 5’ splice site of the PE, respectively (**Fig. 6A**). To enhance the RNA-binding affinity and stability of ASOs, we designed ASOs with two different types of base modifications. The first modification was a uniform phosphorothioate backbone and methoxyethyl at the 2′ ribose position (MOE). The second modification was locked nucleic acid (LNA) modification, where a methylene bridge was introduced linking the 2′ oxygen to the 4′ carbon of the pentose ring. These kinds of modifications were previously described to enhance RNA binding affinity and improve resistance to both nucleases- and RNase H-mediated cleavage of the target RNA (Rigo et al. 2012; Havens and Hastings 2016; Nomakuchi et al. 2016; Ma et al. 2022). We set out our initial screening of ASOs for three different concentrations at lower nanomolar ranges (50 nm, 75 nm, and 100 nm). We first tested ASOs targeting the minigene model. We transfected *EZH2*CCWG (Native) minigene reporter and MOE ASOs (50 nm, 75 nm, and 100 nm) into K562(DOX)SRSF2^P95H^ cells treated with DOX. As control, we used no ASO treatment mock control (CTRL). We isolated RNA after 48 h of transfection and quantified *EZH2*CCWG minigene reporter-specific poison exon inclusion using RT-PCR. In our screening, among the four ASOs, we found ASO2 significantly reduced the poison exon inclusion in *EZH2*CCWG minigene in all tested concentrations, with a better efficiency for a higher concentration (**Fig. 6B**). We also used three additional control MOE ASOs targeting unrelated genes, *HBB, MECP2,* and *MUSK,* respectively. These ASOs showed no effect on *EZH2* poison exon inclusion at any of the tested concentrations (50 nm, 75 nm, and 100 nm); therefore, they rule out any experimental artifacts. We then repeated the experiment with LNA ASOs and observed similar results to those of MOE ASOs (**Fig. 6C**). We also performed these experiments in HeLa(DOX)SRSF2^P95H^ cells treated with DOX. We observed similar results to those of K562(DOX)SRSF2^P95H^. These results suggest that ASOs designed with two different types of chemical modifications possibly function via the same mechanism of action and are not affected by cell types. Note that ASO2 was designed targeting the binding motif 1 of SRSF2^Mut^, which was identified as a crucial motif for exon inclusion. Therefore, the ASO2-mediated splice-switching is highly likely due to blocking the binding of SRSF2^Mut^ at motif 1.

**Figure 6.**
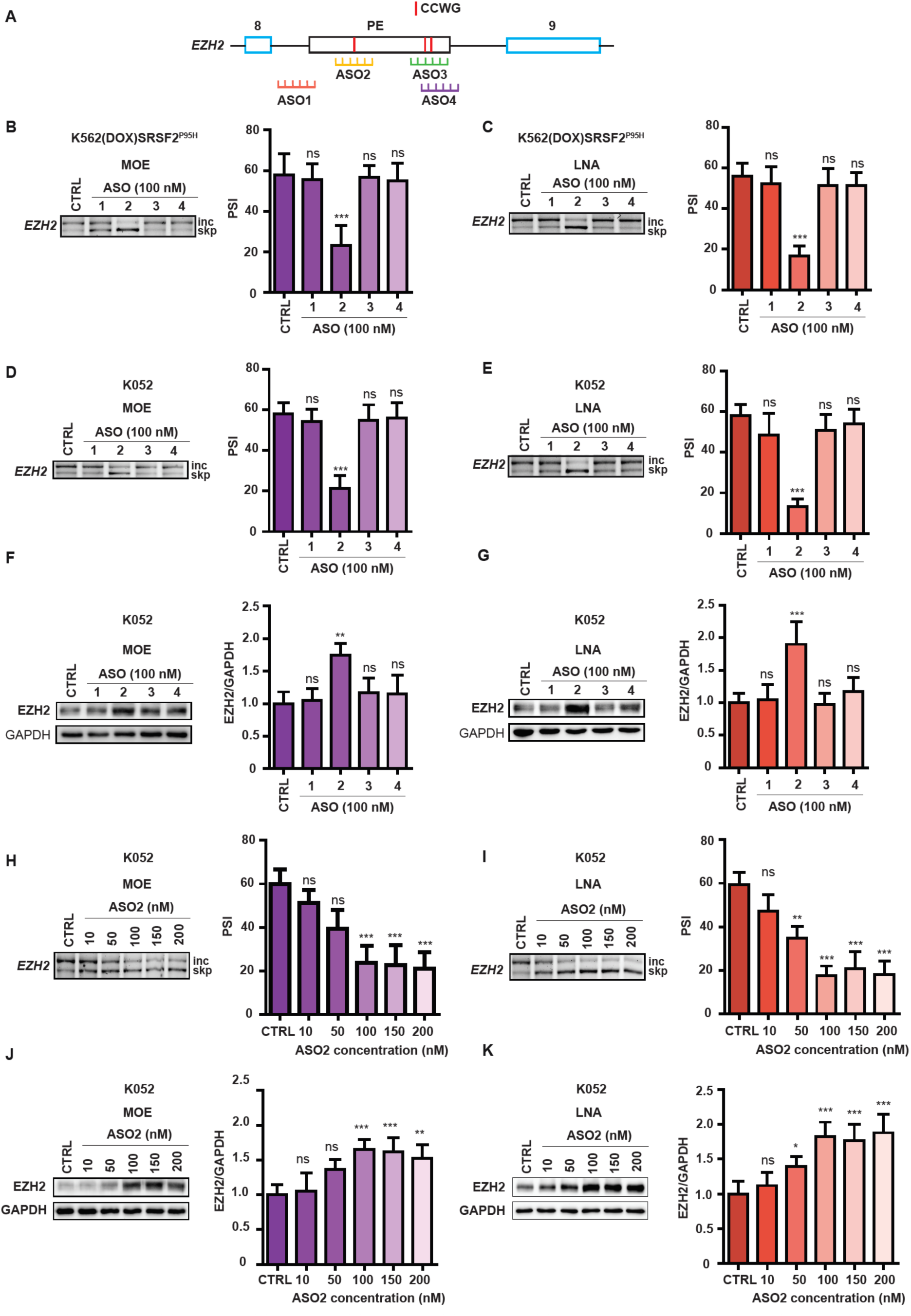
ASO-mediated correction of aberrant poison exon inclusion in *EZH2* in SRSF2-mutated blood cancer. **(A)** Diagram (not to scale) of *EZH2*CCWG minigene reporter and schematics of ASOs and their targeting sites. **(B, C)** Left, representative RT-PCR gel showing *EZH2* minigene reporter-specific poison exon inclusion (inc) and skipping (skp) of transfected K562(DOX)SRSF2^P95H^ cells in the presence of DOX and treated with 100 nM MOE **(B)** or LNA **(C)** ASOs targeting *EZH2*. Right, quantification of *EZH2*CCWG minigene reporter-specific poison exon inclusion as percent spliced-in (PSI) is shown by a bar graph. **(D, E)** Left, representative RT-PCR gel showing endogenous *EZH2* poison exon splicing of K052 cells transfected with 100 nM MOE **(D)** or LNA ASOs **(E)**. Right, quantification of endogenous *EZH2* poison exon inclusion as PSI is shown by a bar graph. **(F, G)** Left, representative western blotting of EZH2 protein expression of K052 cells transfected with 100 nM MOE **(F)** or LNA ASOs **(G)**. GAPDH was used as an internal control. Right, quantification of relative EZH2 protein expression is shown by a bar graph. **(H, I)** Left, representative RT-PCR gel showing endogenous *EZH2* poison exon splicing of K052 cells transfected with the indicated doses of MOE **(H)** or LNA ASO2 **(I)**. Right, quantification of endogenous *EZH2* poison exon inclusion as PSI is shown by a bar graph. **(J, K)** Left, representative western blotting of EZH2 protein expression in K052 cells transfected with the indicated doses of MOE **(J)** or LNA ASO2 **(K)**. GAPDH was used as an internal control. Right, quantification of relative EZH2 protein expression is shown by a bar graph. CTRL: no ASO treatment mock control. All bar graphs (mean ± SD, n=3, *p<0.05, **p<0.01, ***p<0.001, ns: not significant, t test).

### Modulation of *EZH2* poison exon splicing using antisense oligonucleotides in SRSF2^Mut^ cancer model

Since our ultimate goal is to restore the EZH2 protein expression for therapeutic intervention, we next tested the effects of ASOs to correct aberrant splicing of endogenous *EZH2* in SRSF2-mutated cancer cells. Although aberrant poison exon inclusion of *EZH2* was consistently identified in cancer patients with mutated SRSF2, both in RNA-seq data and RT-PCR validation, the efficiency of poison exon inclusion was variable in model cell lines across different studies (Zhang et al. 2015; Rahman et al. 2020b; Kim et al. 2015; Yoshimi et al. 2019; Liang et al. 2018). This disparity could be potentially due to the heterogeneous genetic background in cancer patients compared to engineered cell lines. Therefore, we sought a cell line originating from a cancer patient with an inherent SRSF2 mutation. We finally selected the K052 cell line, originated from a leukemia patient with an inherent SRSF2^P95H^ mutation (as described in **Fig. 1**). Similar to the minigene reporter, we observed that ASO2 (both MOE and LNA) significantly suppressed poison exon inclusion of endogenous *EZH2* (**Fig. 6D and E**) and restored EZH2 protein expression in K052 (**Fig. 6F and G**). To determine whether the extent of ASO2-mediated suppression of *EZH2* poison exon inclusion and restoration of EZH2 protein expression can be titrated, we tested ASO2 (both MOE and LNA) for a dose-response experiment. We transfected ASO2 at increasing concentrations (10 nM, 50 nM, 100 nM, 150 nM, and 200 nM) and harvested cells for both RNA and protein isolation after 48 h of the transfection. We indeed found a dose-dependent effect on splicing correction (**Fig. 6H and I**) and concomitant EZH2 protein expression (**Fig. 6J and K**). These results show the reproducibility and reliability of our experimental screenings (both minigene and endogenous gene) and proof-of-concept.

### Evaluating the on-target specificity of ASO2 in SRSF2^Mut^ cancer model

Since ASOs are short single-stranded nucleic acid molecules, which function by binding to a target RNA, we next evaluated the on-target specificity of our lead ASO (ASO2) in K052 cells. To this end, we checked the splicing profile of several other reported altered splicing events promoted by SRSF2^Mut^, which have been consistently identified in multiple studies. We included additional 11 targets, including *ARMC10, ATF2, CDC45, CDK5RAP2, CRAT, DGUOK, WDR45, INTS3, MELK, SLC25A26, and PEKM* (Zhang et al. 2015; Rahman et al. 2020b; Kim et al. 2015; Yoshimi et al. 2019). We also included K052^WT^ (SRSF2^WT^) as a control to check whether the effect of ASO2 is specific to cancer cells. We found a consistent differential alternative splicing pattern in K052 cells compared to K052^WT^ cells in all 11 targets. Since MOE and LNA ASOs showed consistent results, we included only LNA ASOs for all downstream experiments. Upon treatment of LNA ASO2 in K052 cells, we could not observe any significant changes in alternative splicing in any of the targets, except *EZH2*. These data highlight the on-target specificity of ASO2 only for SRSF2-mutated cancer cells.

### ASO2 enhances methylation of histone H3 at lysine 27 to restore EZH2-mediated chromatin regulation in SRSF2^Mut^ cancer model

We next seek whether ASO2-mediated restoration of EZH2 protein expression rescues EZH2 function. As noted earlier, EZH2 catalyzes methylation of histone H3 at lysine 27 (H3K27me3). We, therefore, analyzed protein levels of histone H3 and H3K27me3 in K052 cells treated with or without LNA ASO2. We found that the overall expression levels of histone H3 remained similar, however, H3K27me3 levels increased significantly upon ASO2 treatment (**Fig. 7A**), suggesting that ASO2 could restore EZH2-mediated chromatin regulation.

**Figure 7.**
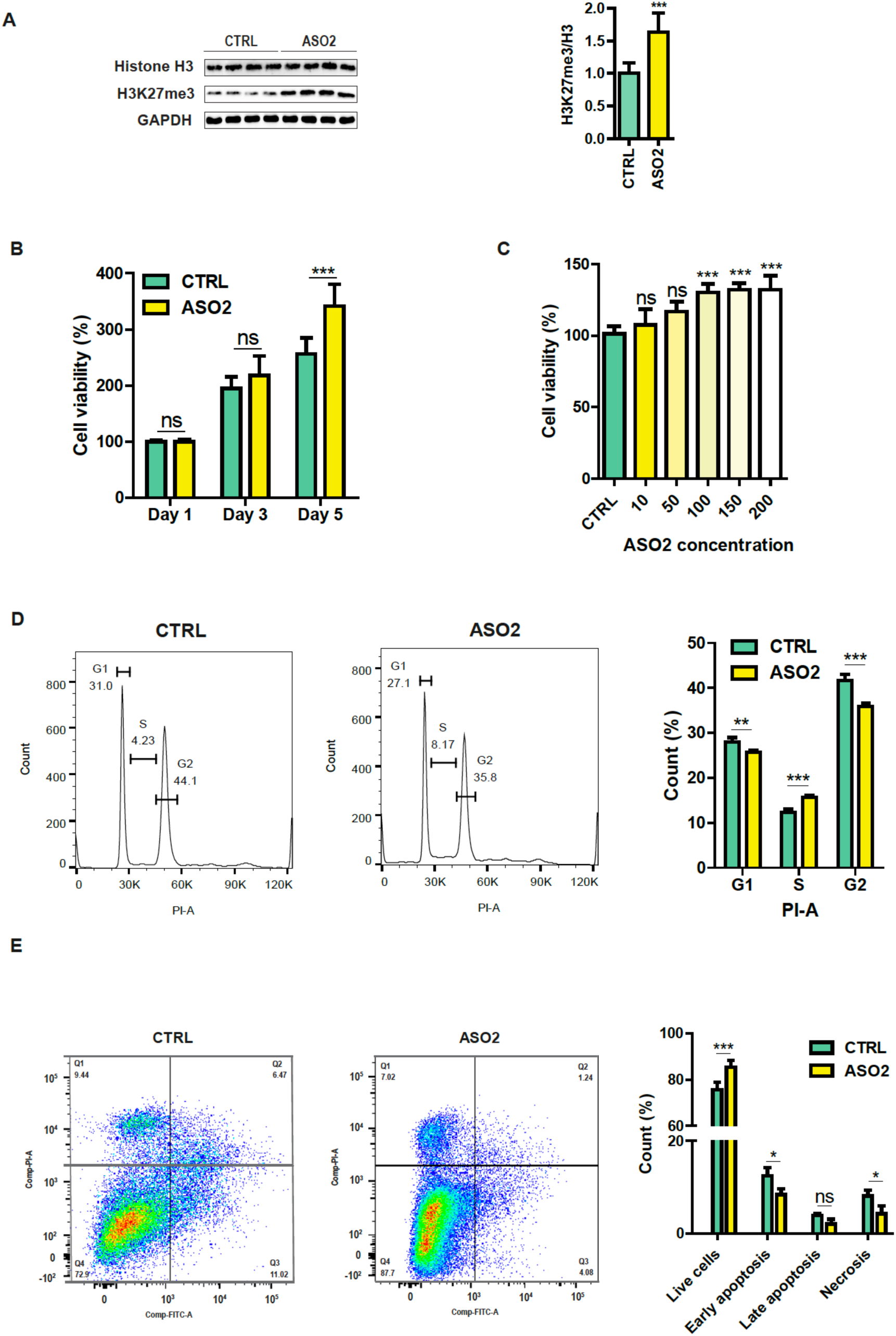
*EZH2* targeting ASO enhances Histone H3K27 methylation, improves cell viability, modulates the cell cycle, and reduces apoptosis. **(A)** Western blotting of Histone H3 and H3K27me3 protein expression in K052 cells treated without (CTRL) or with LNA ASO2 (100 nm) targeting *EZH2*. **(B)** The XTT cell viability assay of K052 in the indicated days treated without (CTRL) or with LNA ASO2 (100 nm). **(C)** The XTT cell viability assay of K052 after 5 days of treatment without (CTRL) or with LNA ASO2 at the indicated doses. **(D)** Cell cycle analysis (left) and relative quantification (right) of K052 after 2 days treated without (CTRL) or with LNA ASO2 (100 nm). **(E)** Apoptosis analysis (left) and relative quantification (right) of K052 after 2 days of treatment without (CTRL) or with LNA ASO2 (100 nm). Bar graphs (mean ± SD, n= 4 for panel (A) and n= 3 for all other panels, *p<0.05, **p<0.01, ***p<0.001, ns: not significant, t test).

### ASO2 improves cell viability, modulates cell cycle kinetics, and reduces apoptosis in SRSF2^Mut^ cancer model

It was previously shown in a murine model that a heterozygous Pro95 mutation in SRSF2 develops MDS with impaired hematopoietic differentiation, altered cell cycle kinetics, and increased apoptosis, resulting in peripheral cytopenia and morphologic dysplasia (Kim et al. 2015). It was further shown that restoring EZH2 expression using a cDNA partially rescues hematopoiesis in *Srsf2*-mutant cells by enhancing cell viability (Kim et al. 2015). We, therefore, tested the effects of LNA ASO2 in rescuing defective hematopoiesis in K052 cells. The XTT cell viability assay revealed that K052 cells treated with 100 nM of LNA ASO2 showed no significant changes in cell viability up to 3 days compared to non-treated cells (**Fig. 7B**). However, after 5 days, cell viability was significantly increased for LNA ASO2-treated cells (**Fig. 7B and C**). We next followed up cell cycle kinetics using flow cytometry. We found that LNA ASO2 treatment reduced the percentage of cells in both G1 and G2 phases, with a concomitant increase in S phase (**Fig. 7D**). In addition, LNA ASO2 treatment reduced apoptosis and necrosis in K052 (**Fig. 7E**). Taken together, these data suggest ASO2 exerts promising effects in rescuing defective hematopoiesis by enhancing cell viability, modulating cell cycle kinetics, and reducing apoptosis and necrosis. Therefore, our lead ASO (ASO2) offers promising potential for clinical development. Our future studies will be focused to evaluate the efficacy of our lead ASO in preclinical animal models *in vivo* and in patient-derived samples.

## Discussion

Since the discovery of recurrent oncogenic mutations in splicing factors, mutations in SRSF2 have been widely studied in hematologic malignancies. Although significant efforts have been employed to identify aberrantly regulated genes, affected pathways, and mechanistic features, the targeted therapeutic development to correct aberrant splicing in SRSF2-mutated cancer has remained limited and ineffective to date. To push the boundary of this underexplored area, we here develop a proof-of-principle to restore the expression of crucial proteins degraded by AS-NMD and linked to tumorigenesis in SRSF2-mutated cancer through an efficient and gene-specific targeted strategy. We scrutinized the poison exon-mediated AS-NMD event in *EZH2*, a crucial epigenetic regulator frequently altered in AML and other cancers (Rahman et al. 2020b; Kim et al. 2015; Yoshimi et al. 2019; Tan et al. 2014; Grimm et al. 2021; Masaki et al. 2019; Sakhdari et al. 2019; Guglielmelli et al. 2011). Our strategy using antisense oligonucleotide pharmacology efficiently corrects aberrant AS-NMD, restores the expression of EZH2 protein, rescues EZH2-mediated chromatin regulation, and improves hematopoiesis and cellular properties.

Although screening of ASOs often exploits random ASOs walking through an entire alternative exon or flanking intronic sequences, we chose to target splicing regulatory regions following precise mechanistic characterization, so that we can develop ASOs with a clear understanding of their mode of action and on-target specificity. AS-NMD is a highly regulated nexus including pre-mRNA splicing in the nucleus, mRNA transport from nucleus to cytoplasm, and mRNA decay in the cytoplasm. Therefore, understanding the complex choreography of regulatory factors in *EZH2* in the entire AS-NMD pathway is highly instrumental in developing efficient ASOs to reverse altered regulation. Since it is well established from previous studies, including ours, that mutations in SRSF2 change its RNA-binding preference from a GGWG motif to CCWG, a logical approach is to develop an ASO to block the binding site of SRSF2^Mut^. However, the CCWG motif is a very short motif, therefore likely to present multiple times throughout a target gene. This represents a primary roadblock to designing an ASO that can cover all binding motifs, since ASOs are short stretches of single-stranded nucleic acid molecules. We therefore focused on identifying a crucial functional motif in *EZH2*. We first focused on three CCWG motifs in *EZH2* PE. Our minigene analyses data clearly demonstrate that the aberrant exon inclusion is regulated due to the presence of an ESE in the PE responsive to SRSF2^Mut^ in SRSF2-mutated cells. In contrast, the absence of an ESE responsive to SRSF2^WT^ in normal cells accounts for the skipping of PE. Systematic and sequential mutagenesis experiments disclosed motif 1 as an essential motif for SRSF2^Mut^-mediated exon inclusion. Artificial recruitment of SRSF2^Mut^ to the specific motif site through MS2 coat protein further confirmed the importance of motif 1. The MS2 hairpin-mediated purification of *EZH2* transcripts (including pre-mRNA and mRNA) and bound RNPs revealed that SRSF2^Mut^ enhances the deposition of several spliceosome components in the flanking splice sites, including U1 snRNP (U1 snRNP 70K, U1 snRNP A, and U1 snRNP C), U2AF65, and U2AF35. These data suggest that the splice sites flanking *EZH2* PE are weakly recognized by the spliceosome, and hence require assistance from a splicing enhancer factor, which we indeed found in our analysis, promoted by SRSF2^Mut^ in SRSF2-mutated cancer.

We also found enhanced deposition of several EJC components and NMD factors in *EZH2* transcripts purified from SRSF2^Mut^ expressing cells compared to SRSF2^WT^ expressing cells. These data from *EZH2* (AS-NMD model) are consistent with our previous data with *HBB* (nonsense mutation-induced NMD model) (Rahman et al. 2020b). We previously revealed that expression of both SRSF2^WT^ and SRSF2^Mut^ is restricted to the nucleus only (Rahman et al. 2020b). The question raised here is how SRSF2^Mut^ augments mRNA decay that happens in the cytoplasm? Note that EJC is a multiprotein complex, consisting of several core and peripheral proteins (Woodward et al. 2017; Singh et al. 2012; Gehring et al. 2009). According to previous studies, EJC is not a preformed complex; rather, it is formed sequentially, with some components assembled in the nucleus and others in the cytoplasm. Furthermore, EJC serves as the anchoring platform for the deposition of NMD factors (UPF proteins) and other mRNA decay factors into an NMD substrate (Woodward et al. 2017; Singh et al. 2012; Gehring et al. 2009). In our data, we found enhanced deposition of core EJC components eIF4A3, Y14, and MAGOH, which are reported to assemble in the nucleus. We didn’t see any enhancement for MLN51. It was suggested that MLN51 may join EJC in the cytoplasm and is not necessary for all transcripts (Gehring et al. 2009; Mabin et al. 2018). Since we found SRSF2^Mut^ expresses only in the nucleus, it can explain the lack of enhancement of MLN51 in *EZH2*. Our data propose that SRSF2^Mut^ enhances the deposition of core EJC components in *EZH2* transcript in the nucleus, which in turn serves as the anchoring platform facilitating the assembly of other peripheral EJC components, NMD factors, and mRNA decay factors, and finally augments mRNA decay in the cytoplasm.

Our mechanistic characterization identified three potential regulatory regions for designing ASOs: ESE motifs in the PE to block the binding of SRSF2^Mut^; flanking 3’ splice site to interfere with the binding of U2AF proteins (U2AF35 and U2AF65); and flanking 5’ splice site to interfere with the binding of U1 snRNP. Therefore, we designed four ASOs targeting these three regions. Among the four, the ASO2 targeting motif 1 site efficiently suppressed PE inclusion and restored EZH2 protein expression. This is in agreement with our mechanistic observation, where motif 1 appeared as a crucial determinant of PE inclusion. To our expectation, ASO3 likely failed as we found that binding motifs 2 and 3 are not sufficient and have minimal additive effects. We speculate that 15-mer ASO1 was not enough to block the binding of U2AF65 and U2AF35 or lacked base pairing affinity to the 3’ splice site. Similarly, ASO4 might fail due to a lack of base pairing affinity. Designing more ASOs surrounding 3’ and 5’ splice sites may identify efficient ASOs. We exploited two different types of base modifications (MOE-PS and LNA) to enhance binding affinity and stability, but both appeared similarly efficient. Therefore, to improve binding affinity, we probably have to consider the base composition and melting temperature (Tm), rather than chemistries.

ASOs could be used differentially to evade NMD. We previously showed that ASO-mediated blocking of EJC binding downstream of a PTC can inhibit SRSF2^Mut^-elicited NMD and can stabilize the PTC-containing transcript (Rahman et al. 2020b). This could be exploited to avoid mRNA degradation and permit protein synthesis. However, a PTC-containing substrate would likely generate a truncated protein, and that would be beneficial only if it preserves essential or functional domains. Therefore, this strategy might not work for many genes. Another strategy could be using ASO along with a translation read-through compound (RTC) (Nomakuchi et al. 2016). In the presence of an RTC, the stop codon is recognized as a triplet coding for an amino acid by the translational machinery (Goldmann et al. 2012). RTC, therefore, can promote the translation of a full-length protein from a PTC-containing mRNA (Goldmann et al. 2012). However, NMD limits the efficacy of RTCs due to the degradation of PTC-containing mRNAs (Nomakuchi et al. 2016). It was shown that ASO-mediated NMD inhibition, along with the read-through compound G418 targeting PTC-containing mRNA, successfully restored the full-length protein with greater efficiency (Nomakuchi et al. 2016). However, this approach could be advantageous for nonsense mutation-induced NMD rather than AS-NMD, since RTC will introduce additional amino acids for AS-NMD mRNA and may interfere with protein structure and function. The best strategy targeting AS-NMD is the splice-switching ASO that we have shown here. Splice-switching ASOs modulate aberrant splicing, evade AS-NMD, and promote expression of the canonical protein.

As noted earlier, in SRSF2-mutated cancer, only a handful of aberrantly spliced targets were identified and validated with a proven link to tumorigenesis (such as *EZH2, INTS3, and CLK3*). ASO correcting one gene will show partial rescue and improvement, and therefore a limitation for cancer therapy compared to monogenic genetic diseases. To address this shortcoming, the combinatorial treatment of ASOs correcting multiple genes will be more beneficial. However, careful consideration should be given to select the overall dose for combinatorial therapy to ensure the safety margin. Another alternative could be using decoy oligonucleotides (spanning several motifs of SRSF2^Mut^ binding sites as a sense strand), which will sequester SRSF2^Mut^ and may prohibit its binding to target genes, therefore, modulating aberrant splicing of all target genes simultaneously (Denichenko et al. 2019). Our future research will extend our proof-of-principle to develop ASOs for combinatorial therapy targeting multiple genes or decoy oligos and will test in preclinical models and hopefully progress to clinical development.

## Materials and methods

### Patient samples

All human related studies were approved by the Institutional Review Boards of the University of Arkansas for Medical Sciences, Memorial Sloan Kettering Cancer Center, Cold Spring Harbor Laboratory, Johns Hopkins University School of Medicine and conducted in accordance with the Declaration of Helsinki protocol. AML patient samples were obtained from de-identified patients at Memorial Sloan Kettering Cancer Center.

### Minigene reporters

The native *EZH2* minigene reporter (*EZH2*CCWG) was kindly shared by Dr. Robert Bradley (Fred Hutchinson Cancer Center) (Kim et al. 2015). We generated different versions of the *EZH2* minigene reporter using site-directed mutagenesis. We confirmed the accuracy of the reporters by sequencing the entire inserts.

### cDNA constructs

T7-SRSF2^WT^, T7-SRSF2^P95H^, MS2, MS2-T7-SRSF2^WT^, and MS2-T7-SRSF2^P95H^ cDNA constructs were previously reported (Rahman et al. 2020b). We generated DOX-inducible FLAG-tagged SRSF2^WT^ and SRSF2^P95H^ cDNA constructs in pInducer20 (Addgene) (Meerbrey et al. 2011).

### Cell culture

K562, K562^Mut^, MOLM-13, SET2, K052, and K052^WT^ cells were cultured in RPMI-1640 medium, whereas HeLa and 293T cells were cultured in DMEM media. All cells were grown with respective medium supplemented with 10% fetal bovine serum (FBS) and maintained in an incubator at 37^°^C with 5% CO_2_. K562(DOX)SRSF2^WT or P95H^ or HeLa(DOX)SRSF2^WT or P95H^ cells were stimulated by 1 µg/ml doxycycline (DOX).

### Cell transfection

Cells were transfected with Lipofectamine 3000 reagent (Invitrogen) according to the manufacturer’s instructions.

### Lentiviral transduction

293T cells were cultured in DMEM medium with 10% FBS with antibiotics. The next day, medium was replaced with fresh DMEM medium with 10% FBS, chloroquine diphosphate, and without antibiotics. Cells were transfected with the plasmid of interest, VSV.G, and psPAX2, and cells were incubated at 37°C for 6 hours. Medium was replaced with fresh DMEM medium with 10% FBS and without antibiotics. After 48 hours, the lentiviral supernatant was collected and filtered through 0.45 mm filters. K562 cells were grown in RPMI-1640 medium with 10% FBS. The next day, K562 cells were transduced with lentiviral supernatant along with polybrene and repeated with fresh virus the following day. We transduced HeLa cells grown in DMEM medium with 10% FBS in a similar way as described for K562 cells.

### Splicing analysis using reverse transcription PCR (RT-PCR)

Total RNA was extracted 48 hours post-transfection of minigene reporter (unless otherwise mentioned) using TRIzol reagent, followed by DNase treatment. Reverse transcription reaction was performed with ImProm-II reverse transcriptase (Promega). During cDNA synthesis, we used oligo-dT reverse primer to check endogenous gene splicing. In contrast, we used a minigene reporter-specific reverse primer during cDNA synthesis to check minigene reporter-specific splicing events. PCR reaction was performed by Phusion High-Fidelity PCR Master Mix with HF Buffer (M0531L, New England Biolabs).

### Western blotting

For protein isolation, cells were washed and harvested in 1x PBS containing Protease Inhibitor Cocktail. Following centrifugation at 2,000 x g for 5 min, the cell pellets were resuspended in buffer A [10 mM HEPES-NaOH (pH 7.8), 10 mM KCl, 0.1 mM EDTA, 1 mM DTT, 0.5 mM PMSF, 0.1% Nonidet P-40, Protease Inhibitor Cocktail], and incubated for 30 min on ice. The samples were then sonicated and centrifuged at 20,000 x g for 5 min. The supernatants were collected as total cell lysate for western blotting.

### MS2-mediated RNA affinity purification

K562(DOX)SRSF2^WT or P95H^ cells were stimulated by 1 µg/ml doxycycline (DOX) in 15 cm plates. After overnight culture, the cells were transfected with *EZH2*CCWG-3xMS2 or *EZH2*AAWG-3xMS2 reporter using Lipofectamine 3000 and then incubated for 48 hours. Cells were then treated with cycloheximide (CHX) at a concentration of 100 µg/ml for two hours before harvesting the cells. The cells were first washed with phosphate-buffered saline (PBS) containing 100 µg/ml CHX, and then resuspended in 3 ml of lysis buffer (20 mM Tris-HCl, pH 7.5, 15 mM NaCl, 10 mM EDTA, 0.5% NP-40, 0.1% Triton X-100, RNase, protease inhibitor cocktail, and 100 µg/ml CHX), followed by incubation on ice for 10 minutes. Cell lysis was done through sonication on ice, and the salt (NaCl) concentration was adjusted to 200 mM. The lysate was clarified by centrifugation at 15,000g for 10 minutes at 4°C, then diluted to a final volume of 10 ml with lysis buffer containing 200 mM NaCl. The lysate was incubated with prewashed amylose resin beads (50% slurry) containing 150 µg of recombinant MBP-MS2 (MS2 coat protein-tagged maltose-binding protein)(Rahman et al. 2020b) for 4 hours at 4°C. The RNA-protein complexes (RNPs) bound to the beads were washed three times with wash buffer (20 mM Tris-HCl, pH 7.5, 200 mM NaCl, 0.1% NP-40) and eluted using elution buffer (20 mM Tris-HCl, pH 7.5, 200 mM NaCl, 10 mM 2-mercaptoethanol, 10 mM maltose, 1 mM PMSF). The purified proteins were subsequently resolved by SDS-PAGE and analyzed by western blotting.

### Antisense oligonucleotides (ASO)-mediated splicing correction

The ASOs were designed targeting the flanking splice sites of the *EZH2* poison exon, or specific binding motifs of SRSF2^Mut^ spanning the poison exon. We synthesized ASOs from Integrated DNA Technologies (IDT) with two different types of modifications on the sugar moiety to prevent RNase H activation. The first type of modification was a uniform phosphorothioate backbone and methoxyethyl at the 2′ ribose position (2′MOE-PS). The second type of modification was locked nucleic acid (LNA) modification, where a methylene bridge was introduced linking the 2′ oxygen to the 4′ carbon of the pentose ring. We transfected ASOs in K562(DOX)SRSF2^P95H^ or KO52 cells using Lipofectamine 3000.

### Analyses of cell viability, and cellular phenotypes

We tested the effects of ASOs on cell viability, and cell cycle kinetics of K052 at different concentrations (10 to 200 nM) in different time periods (1 to 5 days). For cell viability, we evaluated the percentage of K052 cell survival by the XTT Cell Viability Assay (Cat#X12223, ThermoFisher Scientific), and counted cell numbers by QuadCount Automated Cell Counter (Cat#E7500) after ASO treatment.

### Flow cytometry analysis

The ASO-transfected K052 cells and non-treated cells were stained by propidium iodide (PI) (Cat# V13242, ThermoFisher Scientific) for cell cycle and by Annexin V and PI for apoptosis, and then evaluated using BD LSRFortessa-Flow Analyses according to the manufacturer’s instructions. The data were evaluated by FlowJo software, and the results were compared to the control group.

### Quantification and statistical analysis

Densiometric analysis of RT-PCR or western blotting was done by ImageJ software. Statistical analysis was performed either by t-test or one-way ANOVA followed by Tukey’s Multiple Comparison Test. Data are presented as mean and standard deviation, indicated by error bars, and all plots were generated using GraphPad Prism software.

## Data availability

No datasets were generated in this study. We reanalyzed previously published RNA-sequencing data set for AML patients from the Cancer Genome Atlas (TCGA-LAML)(Rahman et al. 2020b). Any additional information related to the data reported in this paper is available from the lead contact upon request.

## Acknowledgments

We are grateful to Robert Bradley (Fred Hutchinson Cancer Center) for sharing the *EZH2* minigene. M.A.R. was supported by the National Institutes of General Medical Sciences (NIGMS) of the National Institutes of Health (R35GM154991 and P20GM152281), Edward P. Evans Foundation, and Winthrop P. Rockefeller Cancer Institute.

## Author contributions

M.A.R. conceived the project. M.A.R. supervised the study. M.A.R., A.R.K., M.R.I., and P.N. designed experiments. M.R.I. and P.N. performed all the experiments with the help of N.M., S.A.H., M.M.H., and S.K.; O.A.-W. provided AML patient RNA samples, human leukemia cell lines, A.R.K. provided MS2-tagged cDNA constructs, and W.B.D. and P.T. provided the K052^WT^ cell line. O.A.-W. and A.R.K. helped in critical data interpretation and shared analytical protocols. M.R.I., P.N., and M.A.R. wrote the primary manuscript. M.A.R., O.A.-W., and A.R.K. finalized the manuscript. All authors approved the final version of the manuscript.

## Competing interest statement

M.A.R., M.R.I., and P.N. are inventors on a patent application related to this study. O.A.-W. is one of the founders and scientific advisors of Codify Therapeutics; he holds equity and receives research funding from this company. O.A.-W. has served as a consultant for Amphista Therapeutics and MagnetBio and is on scientific advisory boards of Envisagenics Inc. and Harmonic Discovery Inc.; O.A.-W. received research funding from Nurix Therapeutics, Minovia Therapeutics, and LOXO Oncology unrelated to this study. A.R.K. discloses the following commercial associations, unrelated to the present work: Stoke Therapeutics (Co-Founder, Director and Chair of SAB); SABs of Skyhawk Therapeutics, Envisagenics, and Autoimmunity BioSolutions; and Consultant for Biogen, SEED Therapeutics, Crucible Therapeutics, Cajal Neuroscience, and Collage Bio.

